# Inferring causal genes at type 2 diabetes GWAS loci through chromosome interactions in islet cells

**DOI:** 10.1101/2022.09.19.508549

**Authors:** Jason M. Torres, Han Sun, Vibe Nylander, Damien J. Downes, Martijn van de Bunt, Mark I. McCarthy, Jim R. Hughes, Anna L. Gloyn

## Abstract

Resolving causal genes for type 2 diabetes at loci implicated by genome-wide association studies (GWAS) requires integrating functional genomic data from relevant cell types. Chromatin features in endocrine cells of the pancreatic islet are particularly informative and recent studies leveraging chromosome conformation capture (3C) with Hi-C based methods have elucidated regulatory mechanisms in human islets. However, these genome-wide approaches are less sensitive and afford lower resolution than methods that target specific loci. To gauge the extent to which targeted 3C further resolves chromatin-mediated regulatory mechanisms at GWAS loci, we generated interaction profiles at 23 loci using next-generation (NG) Capture-C in a human beta cell model (EndoC-βH1) and contrasted these maps with Hi-C maps in EndoC-βH1 cells and human islets and a promoter capture Hi-C map in human islets. We found improvements in assay sensitivity of up to 33-fold and resolved 4.8X more chromatin interactions. At a subset of 18 loci with 25 co-localised GWAS and eQTL signals, NG Capture-C interactions implicated effector transcripts at five additional genetic signals relative to promoter capture Hi-C through physical contact with gene promoters. Therefore, high resolution chromatin interaction profiles at selectively targeted loci can complement genome- and promoter-wide maps.

## Introduction

Most variants implicated by genome-wide association studies (GWAS) are non-coding and are thought to influence type 2 diabetes risk through regulatory effects on gene expression within physiologically relevant cell types. As such, the process of elucidating causal genes (and their corresponding *effector* transcripts) requires integration of functional genomic and molecular epigenomic information in disease relevant cell types and under relevant conditions. Importantly, regulatory effects on gene expression are facilitated by chromatin interactions and causal genes may be identified by their physical contact with enhancer elements encompassing diabetes-associated variants.

Recent studies have employed methods based on chromatin conformation capture (3C) and implicated genes at GWAS loci associated with type 2 diabetes by mapping chromatin structure in human islets and beta cells (Greenwald et al., 2019; Lawlor et al., 2019; Miguel-Escalada et al., 2019; Su et al., 2022). All 3C protocols involve the same core key steps – fixation of chromatin with formaldehyde, restriction enzyme digestion, and re-ligation of restriction fragments (Davies et al., 2017). The resulting “ligation junctions”, which comprise fragments that are co-localised spatially but may be separated linearly by tens to hundreds of kilobases, are sequenced and incorporated into maps of interacting chromatin. Variations in preparing the 3C library and extracting ligation junctions of interest can influence the resolution, sensitivity, and genomic coverage of the resulting chromatin maps, therefore the choice of 3C-based method can markedly alter the detail of chromatin structure information. These differences may affect the inferences made about islet cell biology and the role of T2D associated GWAS variants.

To date, 3C maps in human islets and beta cells have been based on Hi-C approaches that provide genome-wide coverage (and can be enriched for ligation junctions involving promoters), but afford limited detail at individual loci due to prohibitive sequencing requirements and use of low-resolution restriction enzymes (Davies et al., 2017). Alternatively, the NG Capture-C method enables improved resolution and sensitivity at target loci, also referred to as viewpoints or baits, through enrichment from high-resolution 3C libraries (Davies et al., 2017; Hughes et al., 2014).

To assess the extent to which chromatin maps generated from different 3C-based methods impact mechanistic inference at T2D GWAS loci, we performed a systematic evaluation of 27 gene promoters at 23 loci. We performed *next generation* (NG) Capture-C, which involves a double capture procedure that can enrich for captured fragments by up to 1,000,000-fold (Davies et al., 2015), and targeted promoters in the EndoC-βH1 human beta cell line (Ravassard et al., 2011). We also mapped chromatin interactions using sequenced ligation junctions from recent studies that applied Hi-C in EndoC-βH1 cells and human islets, and promoter-capture (pc) Hi-C in human islets (Lawlor et al., 2019; Miguel-Escalada et al., 2019). By comparing these maps with those from NG Capture-C and incorporating GWAS variants co-localised with expression quantitative trait loci (eQTLs) in human islets, we show how distinct chromatin profiles influence the resolution of causal genes for T2D and glycaemic traits.

## Results

We compared chromatin interaction maps for 27 promotors at 23 loci in human EndoC-βH1 cells, derived from NG Capture-C, with previously published Hi-C maps (Lawlor et al., 2019) in EndoC-βH1 cells and human islets, and with a pcHi-C map (Miguel-Escalada et al., 2019) in human islets. These experiments showed marked differences in sensitivity with the NG Capture-C EndoC-βH1 experiment yielding up to ∼27X more ligation junctions than the Hi-C based studies (**Supplemental Table 2**). We assessed how these experimental differences impacted our ability to resolve chromatin interactions. We applied a Bayesian model implemented in peaky (Eijsbouts et al., 2019) to detect fragments showing significant physical interaction with each of the viewpoint (a.k.a. “bait”) fragments encompassing the targeted promoters. Due to the sparsity of per-fragment ligation junction reads, the peaky algorithm failed to converge (and hence unable to perform statistical tests) for six viewpoints in the pcHi-C islet dataset and for all 27 viewpoints in the Hi-C datasets (**Supplemental Table 3**). In contrast, peaky successfully mapped interactions at all viewpoints in the NG capture-C EndoC-βH1 experiment. After merging adjacent fragments with significant interactions, there were 3.6X as many interactions identified by peaky for NG Capture-C than for pcHi-C. Moreover, the median width of significantly interacting chromatin regions was 14.3-fold shorter, indicating a greater ability to fine-map interactions in addition to increased sensitivity (**Supplemental Table 3**).

We also found that interaction peaks resolved by the NG Capture-C experiment were more significantly enriched for islet regulatory features than those gleaned from the pcHi-C experiment (**Supplemental Figure 1**). Enriched islet features included accessible chromatin peaks (Fisher’s exact test odds ratio [OR]=2.17, 95% CI [1.95, 2.40]), H3K27ac ChIP-seq peaks (OR=2.66, 95% CI [2.43, 2.91]) and active promoter (OR=2.85, 95% CI [2.34, 3.45]) and enhancer elements (e.g. type 1 active enhancer, OR=2.07, 95% CI [1.72, 2.46]).

To assess how different chromatin interaction profiles impact mechanistic inference at GWAS loci, we integrated single nucleotide polymorphisms (SNPs) associated with islet gene expression (i.e. eSNPs) and type 2 diabetes and/or glycaemic traits. Of the 27 captured promoters, 21 corresponded to eGenes implicated at 18 loci by 25 pairs of co-localised eSNP and GWAS variants (**Supplemental Table 1**). A total of 12 of these co-localised signals were supported by either NG Capture-C (n=10) or pcHi-C (n=7), with five receiving support from both methods (**Supplemental Table 4**). Included in this set of five was a signal at the *CAMK1D* locus where a genetic association with type 2 diabetes involving SNP rs11257655 is co-localised with an eQTL involving rs11257658 (linkage disequilibrium *r*^*2*^=0.994). The G allele of rs11257658 is associated with decreased human islet expression of *CAMK1D* which encodes calcium/calmodulin-dependent protein kinase 1D (van de Bunt et al., 2015; Viñuela et al., 2020). Both variants, located ∼82 kb upstream of the *CAMK1D* promoter, map to chromatin that physically interacts with the promoter site, thereby corroborating the eQTL (**Figure 1A**). Although the resolution in the pc-HiC study was markedly lower than that for capture-C, the interaction maps from both experiments support *CAMK1D* as the effector gene at this locus.

**Figure 1.**
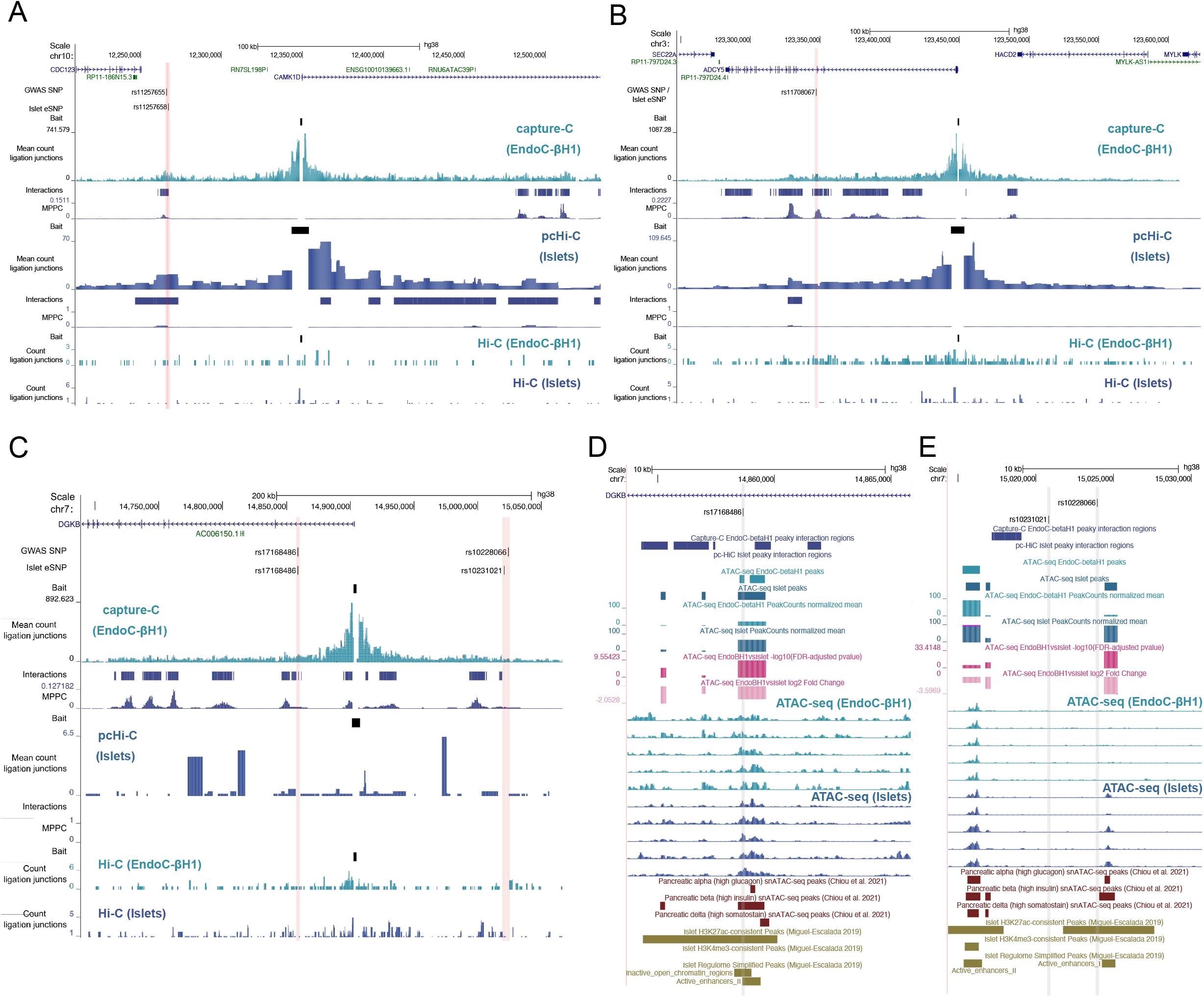
Chromatin interaction profiles at trait-associated loci. Co-localised GWAS-eQTL SNPs are shown with ligation junctions obtained from 3C-based experiments at the **(A)** *CDC123/CAMK1D*, **(B)** *ADCY5*, and **(C)** *DGKB* locus. Tracks of significant chromatin interactions and marginal posterior probability of contact (MPPC) values are shown below the EndoC-βH1 capture-C and islet pcHi-C tracks. Red vertical bars indicate SNP coordinates across 3C-based tracks. Gene annotations correspond to GENCODE V38 protein (blue) and RNA (green) encoding genes. Molecular epigenome profile at the *DGKB* locus is shown for SNPs **(D)** rs17168486 and **(E)** rs10231021 and rs10228066. Differential accessibility between EndoC-βH1 and human islets was assessed using DESeq2 and FDR-adjusted −log10 p-values and log2 fold changes are shown in the dark and light pink, respectively. Select single nuclear ATAC-seq peaks in islet alpha, beta, and delta cells from Chiou et al. 2021 are shown in dark red. Histone post-translational modification ChIP-seq peaks and regulome annotations in human islets from Miguel-Escalada et al. 2019 are shown in dark gold. Grey vertical grey bars indicate SNP coordinates across tracks.

Furthermore, there were five genetic signals supported by chromatin interactions only in our NG Capture-C experiment. This included a co-localised eQTL-GWAS signal fine-mapped to a single SNP (rs11708067) at the *ADCY5* locus where the A allele associates with lower islet expression of *ADCY5*, greater T2D risk, and higher levels of fasting glucose (van de Bunt et al., 2015; Dupuis et al., 2010; Morris et al., 2012; Viñuela et al., 2020; Voight et al., 2010) (**Figure 1B**). We previously reported a chromatin interaction at this locus and allelic imbalance where the risk A allele at rs11708067 associated with decreased chromatin accessibility (Thurner et al., 2018). Another notable example occurred at the *DGKB* locus where there are two independent T2D-associated signals: rs10228066 and rs17168486.

Whereas no significant chromatin interactions were detectable at this locus in the pcHi-C and Hi-C experiments due to low signal, multiple interaction peaks were resolved from the NG Capture-C data, including peaks near the T2D-associated SNPs. The rs17168486 variant, located ∼45 kb upstream of the *DGKB* promoter, mapped within 500 bp of chromatin that significantly interacts with the *DGKB* promoter region (**Figure 1C**). Notably, this SNP – where the T2D risk allele T associates with increased expression of *DGKB* in human islets (Viñuela et al., 2020) – also overlaps enhancer elements in islets and EndoC-βH1 cells (but not in liver, adipose, or skeletal muscle) and accessible chromatin in primary alpha and beta cells (i.e. single-nucleus ATAC-seq peaks) (**Figure 1D**). Moreover, this chromatin accessible region was recently predicted to regulate the expression of *DGKB* in human beta cells with high-insulin content (Chiou et al., 2021), which is further supported by *in vitro* data demonstrating the T2D risk haplotype at rs17168486 influenced luciferase expression in 832/13 and MIN6 cells (Viñuela et al., 2020). In contrast, the rs10228066 variant, located ∼121 kb downstream of the *DGKB* promoter and co-localised with the rs10231021 eQTL signal (LD *r*^*2*^=0.881), mapped more than 1.7 kb from the nearest chromatin interaction. Moreover, neither rs10231021 nor rs10228066 directly overlapped accessible chromatin in EndoC-βH1 or islet cell types (**Figure 1E**). Furthermore, in a recent trans-ethnic GWAS meta-analysis involving 180,834 T2D cases and 1.159M controls, the rs17168486 signal was fine-mapped to a single variant (rs17168486) whereas the other conditionally independent signal at the *DGKB* locus (where the lead SNP rs2215383 is in strong linkage disequilibrium with rs10228066; *r*^*2*^=0.999 in the TOPMED European dataset) was less resolved (13 credible variants and credible interval of 2,318 bp) (Mahajan et al., 2020) and had a credible interval that did not overlap or map within 500 bp of a chromatin interaction. However, a T2D risk haplotype involving variants in high LD with the rs10231021 eSNP did show higher luciferase expression in 832/13 and MIN6 cells with three variants also showing allele-specific binding in a mobility shift assay (Viñuela et al., 2020).

Despite the higher resolution afforded by the NG Capture-C procedure, there were two genetic signals supported by chromatin interactions only in the pcHi-C experiment in human islets: *TCF7L2* and *UBE2E2* (**Supplemental Table 4**). At the *TCF7L2* locus, which has been fine mapped to a single SNP, rs7903146, the T2D risk allele (T) associates with increased *TCF7L2* expression in islets (Viñuela et al., 2020) and the SNP overlaps an islet enhancer element and accessible chromatin in bulk islet tissue and islet alpha, beta, and delta cells (**Supplemental Figure 2, Supplemental Table 5**). Notably, chromatin accessibility at this region was considerably lower in EndoC-βH1 cells than in islets (log2FC=-2.07; FDR-adjusted p-value = 1.68e-07). Therefore, the lower accessibility in this cell-type may explain, in part, the lack of pronounced chromatin interaction at this site from the NG Capture-C profile. In the case of *UBE2E2*, a T2D-associated SNP rs35352848 is co-localised with an eQTL (rs13094957) for *UBE2E2* expression in islets and overlapped a broad islet pcHi-C chromatin interaction with the *UBE2E2* promoter. However, neither variant directly maps to H3K27ac peaks, enhancer elements, or accessible chromatin in islets, or in snATAC-seq peaks in beta, alpha, or delta cells (**Supplemental Table 5**). Notably, a recent trans-ancestry GWAS meta-analysis fine-mapped this signal to six credible variants and a wide credible interval of nearly 200 kb (overlapping 12 chromatin interactions). Therefore, more investigation is needed to resolve the causal variant at this signal.

## Discussion

We found that NG Capture-C achieved substantially greater resolution and sensitivity over Hi-C based approaches at the loci we investigated in human beta cells. This corresponded to enhanced fine-mapping of chromatin interactions that were more enriched for accessible chromatin peaks and islet regulatory elements than interactions gleaned from the pcHi-C experiment. Chromatin interactions provided support for genes implicated by GWAS and eQTL co-localization at 12 of 25 evaluated signals, with five signals supported by interactions in the both the pcHi-C and NG Capture-C experiments. Relative to the pcHi-C study, interactions from our NG Capture-C study corroborated five additional target genes: *ADCY5, DGKB*, and three genes at the *GPSM1* locus (*DNLZ, CARD9*, and *GPSM1*). Experimental studies have implicated both *ADCY5* and *DGKB* in insulin secretion, with *ADCY5* shown to be indispensable for glucose-induced insulin secretion in human islets (Hodson et al., 2014; Peiris et al., 2018). Furthermore, our results corroborate co-localized GWAS and islet eQTL signals at the *GPSM1* locus. Notably, of the signals evaluated in this study, only chromatin encompassing the SNP rs28505901 at this locus showed a significantly stronger interaction in EndoC-βH1 cells relative to LCLs, specifically with the *DNLZ* promoter (**Supplemental Table 4**).

Chromatin interactions from the EndoC-βH1 NG Capture-C experiment did not corroborate candidate disease genes at all evaluated loci, with *TCF7L2* being the most salient exception. We observed lower chromatin accessibility in EndoC-βH1 cells at this site which may reflect an epigenomic profile corresponding to an earlier developmental stage. Hence, the discrepancy at this locus may be a consequence of EndoC-βH1 cells being a fetal-derived cell line rather than evidence against a mechanism of action in beta cells. Due to the relatively modest number of viewpoints interrogated in this study, it is unclear if such discordances are primarily due to inherent epigenomic differences between cell and tissue types. The unavailability of pcHi-C data in EndoC-βH1 cells and NG Capture-C data in islets also limited our comparisons as we could not control for cell-type in our evaluation of these two approaches. It is possible that interactions detected in islets but not in EndoC-βH1 cells may reflect enhancer-promoter loops specific to other islets cell types. Notably, a recent study of Hi-C maps in FACS sorted islet cells implicated alpha cells at a T2D-associated signal at the *WFS1* locus, and acinar cells at a signal mapping to the *CPA4* locus (Su et al., 2022). Differences in interaction profiles between islets and EndoC-βH1 cells may also reflect distinct epigenomic features resulting from SV40LT transduction. Additional chromatin maps will be needed to fully address these questions and recent improvements in NG Capture-C technology may make application in rarer cell populations more tractable (Downes et al., 2021). However, we have demonstrated that markedly enriching 3C libraries for promoters of interest can reveal additional interactions at type 2 diabetes and glycaemic trait-associated loci. Therefore, selective capture of fine-mapped genetic loci may greatly complement genome- or promoter-wide chromatin maps.

## Author contributions

J.M.T, V.N, M.I.M, and A.L.G conceived and planned the main analysis. J.M.T. conducted bioinformatic and statistical analysis of the NG Capture-C sequencing data. H.S. performed bioinformatic analyses of the Hi-C and pcHi-C sequencing data. V.N. and D.J.D. cultured EndoC-βH1and LCL cells and performed the NG Capture-C protocol. M.v.d.B. contributed to the experimental design and selection of loci to target for NG Capture-C. J.R.H. and D.J.D. provided valuable insight into the analysis and interpretation of the NG Capture-C experiment. M.I.M. and A.L.G. supervised this work. J.M.T. wrote the first draft of the manuscript and J.M.T. and A.L.G. edited the manuscript. All authors approved the manuscript.

## Acknowledgments

A.L.G. is a Wellcome Senior Fellow in Basic Biomedical Science. A.L.G. is funded by the Wellcome (200837) and National Institute of Diabetes and Digestive and Kidney Diseases (NIDDK) (U01-DK105535; U01-DK085545, UM1DK126185). This work was carried out as part of the WIGWAM Consortium (Wellcome Investigation of Genome Wide Association Mechanisms) funded by a Wellcome Trust Strategic Award (106130/Z/14/Z) to M.I.M, J.H and A.L.G. This research was also supported by the Wellcome Trust Core Award Grant Number 203141/Z/16/Z with additional support from the NIHR Oxford BRC. The views expressed are those of the author(s) and not necessarily those of the NHS, the NIHR or the Department of Health.

## STAR Methods

### RESOURCE AVAILABILITY

#### Lead contact

Further information and requests for resources and reagents should be directed to and will be fulfilled by the lead contact, Jason Torres.

#### Materials availability

This study did not generate new unique reagents.

#### Data and code availability

NG capture-C sequencing data from EndoC-βH1 cells and LCL cells have been deposited on the European Genome-phenome Archive (EGA), which is hosted by the European Bioinformatics Institute of the European Molecular Biology Laboratory (EMBL-EBI) and the Centre for Genomic Regulation (CRG), under accession number EGAS00001006105 and will be released upon publication. ATAC-seq data from EndoC-βH1 have also been deposited on EGA under accession number EGAS00001006105. Further information about EGA can be found on https://ega-archive.org. All original code has been deposited at Zenodo and is publicly available as of the date of publication. DOIs are listed in the key resources table. This paper analyzes existing, publicly available data. Source data and publicly available resources used for this study supporting all findings are detailed in the key resources table. Any additional information required to reanalyze the data reported in this paper is available from the lead contact upon request.

### EXPERIMENTAL MODEL AND SUBJECT DETAILS

EndoC-βH1 cells (RRID: CVCL_L909) were purchased from Endocell, and cultured as described previously (Hastoy et al., 2018; Ravassard et al., 2011). Lymphoblastoid cell lines (GM12878; RRID: CVCL_7526) were procured from the Coriell Institute. Publicly available *Mbo*I enzyme-based Hi-C sequencing data corresponding to EndoC-βH1 cells and human islets (n=1), were accessed (Lawlor et al., 2019). Promoter capture Hi-C (pcHi-C) data corresponding to human islets from four donors were downloaded from the EGA database (accession EGAS00001002917).

## METHOD DETAILS

### Next Generation Capture-C

Promoters for 27 gene transcripts at 23 loci were selected for capture. These included 21 genes at 18 loci harbouring both islet eQTLs and genome-wide significant associations with type 2 diabetes and/or glycemic traits (van de Bunt et al., 2015; Viñuela et al., 2020). The six remaining genes included three control genes with high expression in lymphoblastoid cell lines (LCLs), the *GCK* gene encoding glucokinase (implicated in monogenic forms of diabetes and hyperinsulinemia), and two genes (*CDKAL1* and *SOX4*) at the *CDKAL1* locus associated with T2D (**Supplemental Table 1**). 70-mer biotinylated oligonucleotide probes (IDT xGen Lockdown oligonucleotides) targeting *Dpn*II restriction fragments were designed using CapSequm (Hughes et al., 2014) with filtering for repetitive elements, duplicates (≤ 2), BLAT density score (≤40), and GC content (≤%60) (Downes et al., 2022).

*In situ* 3C libraries were generated in EndoC-βH1 cells and LCLs by *Dpn*II digestion and T7 ligation chromatin (Davies et al., 2015); 3C material was sonicated to 200 base pairs (bp) and index using NEB Next DNA library prep reagents. Indexed libraries were pooled and double capture performed with Nimblegen SeqCap EZ reagents (Roche) (Davies et al., 2015). Sequencing was performed on the Illumina NextSeq platform with 150 bp paired-end reads. Sequenced reads were mapped to GRCh38 with bowtie using CCseqBasicS (Telenius et al., 2020) which trims adaptor sequences, reconstructs read pairs with flash, conducts an *in silico* digestion of *Dpn*II fragments, maps reads, identifies paired “capture” and “reporter” fragments, and filters PCR duplicates.

### Hi-C and Promoter capture Hi-C

Hi-C sequencing data for EndoC-βH1 cells and human islets were processed using the Juicer (v1.75) pipeline (Durand et al., 2016). Sequencing reads from the *Hind*III-digested pcHi-C library were processed and mapped to genome build GRCh38 using HiCUP (v0.8.1) (Wingett et al., 2015). Bait and prey fragments were quantified using chicagoTools from the CHiCAGO package (v1.14.0) (Cairns et al., 2016). Promoter bait design coordinates for genome build hg19 were obtained from this study (Javierre et al., 2016). and lifted over to GRCh38 using LiftOver.

### ATAC-seq

ATAC library preparation and sequencing was performed for 11 EndoC-βH1 cell lines using the same protocols used in Thurner et al. 2018 (Thurner et al., 2018). Human islet ATAC sequencing data corresponding to 13 donors from the Miguel-Escalada et al. 2019 study was accessed from the EGA database (accession EGAS00001002917). Sequencing reads were processed and mapped to GRCh38 using the ENCODE ATAC-seq bioinformatic pipeline (v1.9.3), with peak calling performed with MACS2 (v2.7.1) (Feng et al., 2012) using default parameters. Reads within peaks were quantified with featureCounts (v2.0.1) (Liao et al., 2019) and normalized by median of observed count ratio *size factors* with DESeq2 (v1.26.0) (Love et al., 2014).

## QUANTIFICATION AND STATISTICAL ANALYSIS

NG Capture-C reporter counts for each replicate (n=3) of EndoC-βH1 and LCL cells were normalized to the number of *cis* reporter counts (i.e. same chromosome) per 100,000 *cis* reporter reads with CaptureCompare (Telenius et al., 2020). Chromatin interaction mapping in NG Capture-C, pcHi-C, and Hi-C datasets was performed with peaky (Eijsbouts et al., 2019) using recommended settings (omega = −3.8). Interactions were considered significant if the marginal posterior probability of contact (MPPC) exceeding 0.01 within a range of 250 kb to the viewpoint (i.e. captured fragment) or 0.1 between 250 kb and 1 Mb relative to the viewpoint. Differential chromatin interactions (NG capture-C) between EndoC-βH1 and LCL cells, and differentially accessible (ATAC-seq) peaks between EndoC-βH1 and primary human islet cells were called using DESEq2 (v1.26.0).

## Supplemental item titles and legends

**Supplemental Table 1: Description of captured loci**. Genes targeted in the next-generation capture-C experiment, and evaluated in the promoter capture Hi-C and Hi-C experiments, are listed with their corresponding loci, rationale for inclusion, known GWAS associations with T2D and/or glycemic traits, and expression quantitative trait loci (eQTL) SNPs (i.e. eSNPs) associated with gene expression in primary human islets. GWAS-eQTL colocalization analysis results reported by the Integrated Network for Systematic analysis of Pancreatic Islet RNA Expression (InsPIRE) consortium are also included from Viñuela et al. 2020.

**Supplemental Table 2: Comparison of ligation junctions across methods**. For each captured gene, the targeted viewpoint fragment encompassing the gene transcription start site (TSS) is indicated along with descriptive statistics corresponding to the mapped ligation junction reads from the next-generation capture-C experiment in EndoC-betaH1 cells, promoter capture Hi-C experiment in human islets from Miguel-Escalada et al. 2019, and Hi-C experiments in EndoC-betaH1 cells and islets from Lawlor et al. 2019. The overlap region indicates if the TSS overlapped the viewpoint fragment directly (i.e. “viewpoint”) or mapped within 1kb of the viewpoint fragment (i.e. “exclusion”).

**Supplemental Table 3: Comparison of significant chromatin interactions mapped with peaky**. For each captured locus, the number of restriction fragments with marginal posterior probability (MPPC) exceeding 0.01 within 250k of the viewpoint fragment (i.e. proximal interaction) or exceeding 0.10 within 1Mb of the viewpoint fragment (i.e. distal interaction) are shown for the capture-C and pcHi-C experiments. Interaction peaks are delineated by merging adjacent fragments meeting MPPC significance thresholds.

**Supplemental Table 4: Integration of peaky interactions with GWAS variants and eSNPs**. Significant chromatin interactions at restriction fragments encompassing reported T2D and/or glycemic trait associated SNPs and/or SNPs associated with gene expression in human islets are shown for the next generation capture-C and pcHi-C experiments. GWAS and eSNPs reported to be significantly colocalized in Viñuela et al. 2020 are indicated. For the capture-C experiment, chromatin that interacted more strongly in EndoC-betaH1 cells relative to LCL cells is also indicated.

**Supplemental Table 5: Single cell accessible chromatin in pancreas at captured loci**. T2D/glycemic trait associated SNPs and islet eSNPs at evaluated loci overlapping single nuclear ATAC-seq peaks in pancreas cell types reported in Chou et al. 2021 are indicated.

