## Supplemental figures for "Inferring causal genes at type 2 diabetes GWAS loci through chromosome interactions in islet cells"

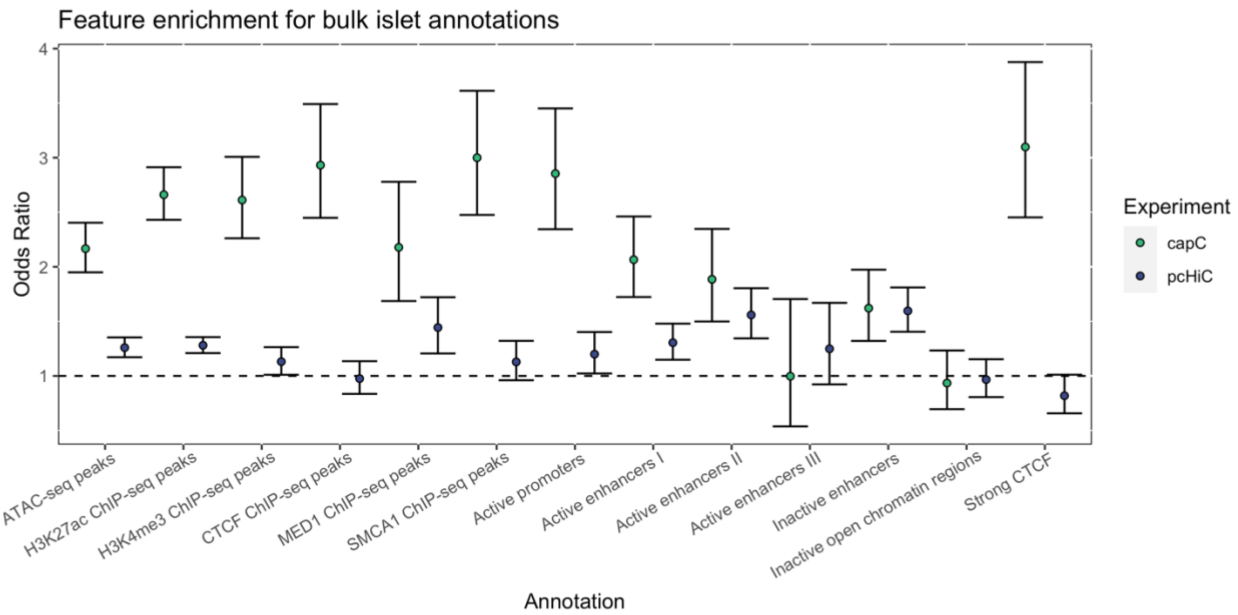


**Supplementary Figure 1. Enrichment of islet epigenomic features at regions with chromatin interactions.** Chromatin interactions with the captured promoters from the EndoC-βH1 capture-C (green) and islet pcHi-C (blue) experiments were mapped with peaky and evaluated for enrichment of islet ATAC-seq, histone ChIP-seq, and regulome features from Miguel-Escalada et al. 2019. Enrichment across all captures was assessed by binning 1kb segments within 1Mb of each targeted promoter and performing Fisher's exact test for each set of islet features and chromatin interactions. The three negative control viewpoints (*CR2*, *PAX5*, and *TNFSF11*) were excluded from enrichment analysis.

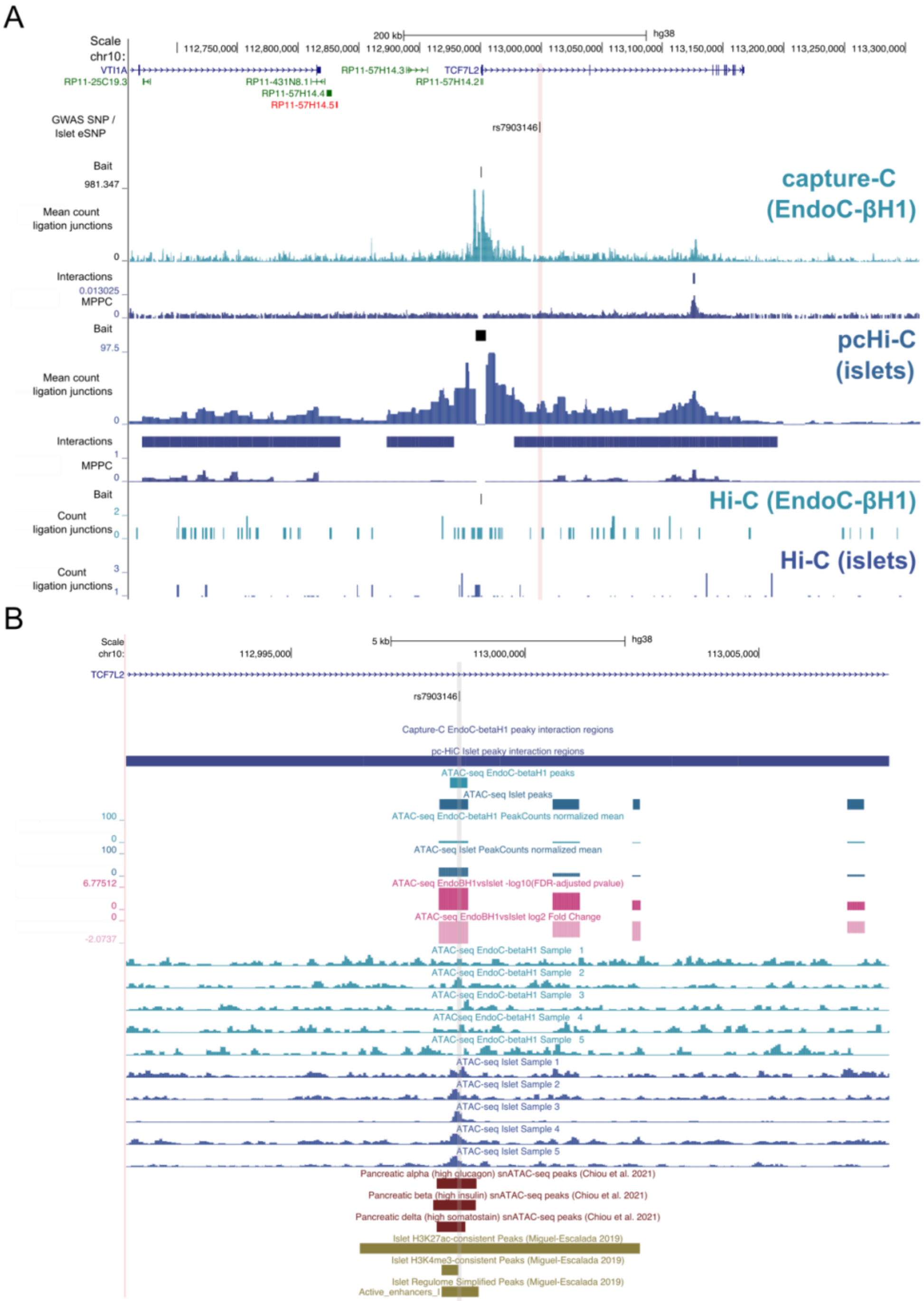


**Supplementary Figure 2. Molecular epigenome profile at the TCF7L2 locus. (A)** Co-localised GWAS-eQTL SNPs are shown with ligation junctions obtained from 3C-based experiments. Tracks of significant chromatin interactions and marginal posterior probability of contact (MPPC) values are shown below the EndoC-βH1 capture-C and islet pcHi-C tracks. Red vertical bars indicate SNP coordinates across 3C-based tracks. Gene annotations correspond to GENCODE V38 protein (blue) and RNA (green) encoding genes. Maps of chromatin accessibility in EndoC-βH1 and human islets are shown for **(B)** rs7903146. Differential accessibility between EndoC-βH1 and human islets was assessed using DESeq2 and FDR-adjusted -log10 p-values and log2 fold changes are shown in the dark and light pink, respectively. Select single nuclear ATAC-seq peaks in islet alpha, beta, and delta cells from Chiou et al. 2021 are shown in dark red. Histone post-translational modification ChIP-seq peaks and regulome annotations in human islets from Miguel-Escalada et al. 2019 are shown in dark gold. Grey vertical grey bars indicate SNP coordinates across tracks.
